# SCOUR: A stepwise machine learning framework for predicting metabolite-dependent regulatory interactions

**DOI:** 10.1101/2021.05.14.444159

**Authors:** Justin Y. Lee, Britney Nguyen, Carlos Orosco, Mark P. Styczynski

**Author notes:** Corresponding author (MPS).

## Abstract

**Background:** The topology of metabolic networks is both well-studied and remarkably well-conserved across many species. The regulation of these networks, however, is much more poorly characterized, though it is known to be divergent across organisms – two characteristics that make it difficult to model metabolic networks accurately. While many computational methods have been built to unravel transcriptional regulation, there have been few approaches developed for systems-scale analysis and study of metabolic regulation. Here, we present a stepwise machine learning framework that applies established algorithms to identify regulatory interactions in metabolic systems based on metabolic data: Stepwise Classification Of Unknown Regulation, or SCOUR.

**Results:** We evaluated our framework on both noiseless and noisy data, using several models of varying sizes and topologies to show that our approach is generalizable. We found that, when testing on data under the most realistic conditions (low sampling frequency and high noise), SCOUR could identify reaction fluxes controlled only by the concentration of a single metabolite (its primary substrate) with high accuracy. The positive predictive value (PPV) for identifying reactions controlled by the concentration of two metabolites ranged from 32-88% for noiseless data, 9.2-49% for either low sampling frequency/low noise or high sampling frequency/high noise data, and 6.6-27% for low sampling frequency/high noise data, with results typically sufficiently high for lab validation to be a practical endeavor. While the PPVs for reactions controlled by three metabolites were lower, they were still in most cases significantly better than random classification.

**Conclusions:** SCOUR uses a novel approach to synthetically generate the training data needed to identify regulators of reaction fluxes in a given metabolic system, enabling metabolomics and fluxomics data to be leveraged for regulatory structure inference. By identifying and triaging the most likely candidate regulatory interactions, SCOUR can drastically reduce the amount of time needed to identify and experimentally validate metabolic regulatory interactions. As high-throughput experimental methods for testing these interactions are further developed, SCOUR will provide critical impact in the development of predictive metabolic models in new organisms and pathways.

## Background

Biochemists have amassed a large amount of knowledge about the topology of the chemical reaction network that cells use to transform nutrients into energy and the building blocks for more cells, collectively known as “metabolism”. The substrates, products, and cofactors for hundreds of reactions have been elucidated, from the most central pathways like glycolysis to more distant pathways for the biosynthesis of uncommon metabolites. Many of these pathways are extremely well-conserved across the tree of life [1], with the basics of central carbon metabolism being quite similar from bacteria to humans. What varies much more greatly across species, and what allows such diverse metabolic phenotypes to arise from such otherwise similar reaction networks, is the regulation and utilization of the reactions in those networks. However, this regulation, despite its major importance in the function and diversity of life, is nowhere near as well understood as the topology of the metabolic network [2]. This is especially true for the direct regulation of reactions by metabolites, which is particularly poorly characterized compared to some other levels of regulation like transcriptional regulation. This is in large part due to the difficulty in experimental characterization of direct regulation by metabolites.

One critical form of direct regulation of metabolic reactions (and arguably the most common) is allosteric regulation, where a regulator and a protein (in this case an enzyme) interact at a location other than the active site [3]. In this mechanism, a metabolite that is not the primary substrate of an enzyme binds to that enzyme and inhibits or promotes the reaction rate, most typically via an induced change in protein conformation. While metabolite levels can affect processes on the genome, transcriptome, and proteome levels [4], metabolite-dependent regulation of enzyme reaction rates is extremely important because it results in the control of reactions on a short timescale (less than 30 seconds) due to the direct interaction between metabolite and enzyme rather than requiring intermediate steps like transcription to effect changes [5, 6]. Their prevalence in metabolic systems makes it vital to account for these regulatory interactions to create accurate metabolic models.

Metabolic models that use only the known stoichiometry of the system and exclude metabolite-dependent regulation often have extremely limited accuracy. Machado et al. showed that including allosteric regulation in a model of *E. coli* is vital for predicting flux dynamics and can reveal “metabolic hubs,” where a metabolite is connected to many reactions instead of only the few found in the stoichiometric topology [6]. Despite its prevalence and importance, the exact structure of this regulatory network (which metabolites regulate which fluxes) is typically unknown in all but the best-studied metabolic pathways in the best-studied organisms. With hundreds of metabolites and hundreds of fluxes in any given metabolic network and no effective high-throughput methods for finding metabolite-protein interactions (compared to, for example, protein-protein interactions [7, 8]), the space of possible regulatory interactions is too vast to experimentally explore [9].

In contrast, the comparatively more mature fields of genomics and transcriptomics have yielded a variety of experimental approaches to explore transcriptional regulation, providing sufficient data to motivate the development of computational approaches to map out gene regulatory networks [10, 11]. Some of the latest computational approaches have leveraged spatial and temporal gene expression data, environmental data, and rapid gene perturbation data to better predict and understand gene regulation [12–14]. Machine learning has been used with increasing popularity in recent years to build gene regulatory networks [15–17]. Other efforts have even integrated metabolomics data with transcriptional data to discover what metabolites control transcriptional responses and gene expression [18, 19].

There have been far fewer computational approaches to identify metabolite-dependent regulatory interactions of enzymes in metabolic systems. Link et al. [5] used dynamic metabolite data to fit an ensemble of kinetic models with different putative regulatory interactions to rank which interactions contributed the most to fitting accuracy. Another approach by Hackett et al., named SIMMER [20], estimated kinetic parameters using non-linear optimization to establish if all reactions in a system could be sufficiently explained by Michaelis-Menten kinetics or if additional allosteric parameters were required. While these computational approaches are invaluable in saving time and costs for laboratory experiments, both methods rely on sampling [5] or estimating [20] kinetic parameters, which can be computationally taxing. An approach for identifying metabolite-dependent regulatory interactions without requiring kinetic parameters would be extremely useful for systems biology modeling. Although approaches using protein docking, such as AlloFinder [21], are promising for future *ab initio* prediction of regulatory interactions, current limitations in the accuracy of molecular simulations make systems-scale exploration of allosteric interaction space challenging and motivate a desire for approaches that can exploit increasingly widely available experimental datasets for these purposes.

Here, we present a new machine learning approach for Stepwise Classification Of Unknown Regulation (SCOUR) that leverages metabolomics and fluxomics data to predict likely metabolite-dependent regulatory interactions. SCOUR uses established machine learning algorithms in a stepwise process that focuses on identifying reactions controlled by one, two, or three metabolites. While SCOUR benefits from stepwise, serial inference of these increasingly complex interactions, each step is independent, uses different classification features, and can be performed without the others. Importantly, the classification task that SCOUR looks to address typically has insufficient training data available to generate useful models, so we devise a strategy we refer to as “autogeneration” that we use to create sufficient data to train the models. We test our framework on two synthetic model networks, as well as on models of *S. cerevisiae* and *E. coli* metabolism, to show that SCOUR can be used on a variety of systems. Applying SCOUR to poorly-studied organisms has the potential to enable discovery of previously unknown regulatory interactions that are key to developing accurate and predictive metabolic models.

## Methods

### Metabolic Networks

In this work, we examined four metabolic networks of varying size and complexity: two synthetic model networks and two biological systems. We simulated each metabolic network with fifteen sets of randomly generated metabolite concentration initial conditions (except for the first set of initial conditions for the biological systems, which were kept at their original values) to produce fifteen sets of metabolite concentration and flux data used in the testing sets of SCOUR. Each metabolic network is described in detail below.

### Synthetic Model Networks

To initially test and evaluate SCOUR, we created two small synthetic model networks. The Smaller Synthetic Model (Fig. 1A) contains six metabolites and six reactions, while the Bigger Synthetic Model (Fig. 1B) contains ten metabolites and ten fluxes. Synthetic systems of these sizes are small and simple enough to easily assess the performance of SCOUR while developing the framework, but large enough to emulate biological behavior of metabolite-flux interactions in metabolic systems. In both models, the influx (v_1_) is a constant flux that is not controlled by any metabolites and is not considered when using SCOUR. Both models contain reactions controlled by one, two, or three metabolites, including both positive and negative regulatory interactions. Table 1 summarizes the number of each type of interaction in each model. The network dynamics were defined using Biochemical Systems Theory (BST) equations using power law kinetics for reaction rates [22], with mass action parameters randomly assigned between 0.1 and 1 and regulation parameters randomly assigned between 0.1 and 1 for positive regulatory interactions and between −1 and −0.1 for negative regulatory interactions.

**Fig. 1.**
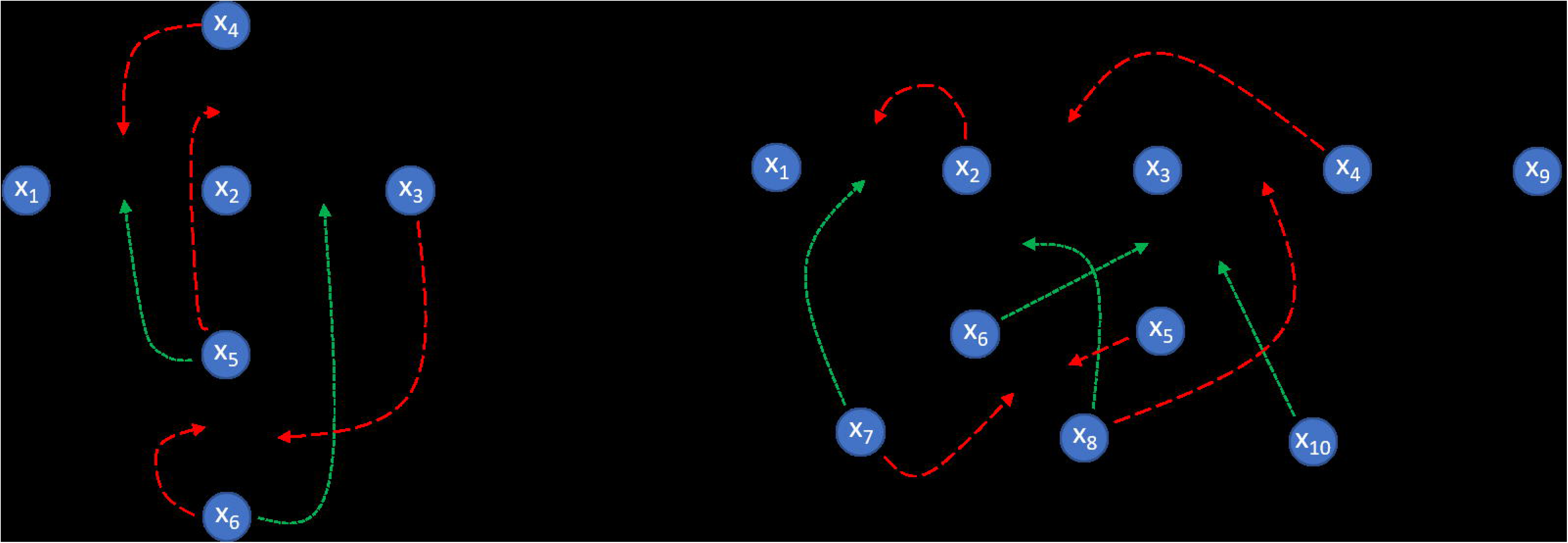
Synthetic Model Networks. Two synthetic model networks created using BST frameworks to generate *in silico* metabolomics and fluxomics data. x_i_ represent metabolites, v_i_ represent reaction fluxes (solid black lines), long-dashed red lines represent regulatory behavior that causes inhibition, and short-dashed green lines represent regulatory behavior that increases activity.

**Table 1:** The number of *n*-controller metabolite interactions that exist in or are possible for each model. For possible interactions, the first number assumes that regulatory interactions are correctly identified at each step of the framework and are removed from consideration for higher-order interactions. The number in parentheses is the total number of possible interactions if a stepwise framework were not used, illustrating the significant decrease in the number of interactions to be assessed in a stepwise framework.

|  | Smaller Synthetic Model | Bigger Synthetic Model | *S. cerevisiae* | *E. coli* |
| --- | --- | --- | --- | --- |
| # of 1-controller interactions | 1 | 3 | 5 | 25 |
| # of possible 1-controller interactions | 5 | 9 | 12 | 39 |
| # of 2-controller interactions | 2 | 3 | 10 | 13 |
| # of possible 2-controller interactions | 20 (25) | 54 (81) | 157 (262) | 243 (668) |
| # of 3-controller interactions | 2 | 3 | 5 | 2 |
| # of possible 3-controller interactions | 20 (50) | 108 (324) | 520 (2720) | 336 (5384) |
| # of 4-controller interactions | N/A | N/A | 2 | 4 |
| # of possible 4-controller interactions | N/A | N/A | 380 (17860) | 480 (27120) |

### Biological Models

To test SCOUR on more biologically relevant systems, we examined a model of central carbon metabolism in *Escherichia coli* and a model of glycolysis in *Saccharomyces cerevisiae* [23, 24]. The *E. coli* model contains 18 metabolites and 48 reactions, while the *S. cerevisiae* model contains 22 metabolites and 24 reactions. We used the previously published kinetic equations and parameters for these systems; in both cases, the mathematical forms of the rate expressions include Michaelis-Menten, Hill, and mass action kinetics. In the *E. coli* model, the fluxes for glucose kinetics, murein synthesis, tryptophan synthesis, and methionine synthesis are constant and not controlled by any metabolites. Likewise, in the *S. cerevisiae* model, the fluxes for glucose mixed flow to extracellular medium and cyanide flow are constant and not controlled by any metabolites. As in the synthetic models, both the *S. cerevisiae* and *E. coli* models include reactions controlled by one, two or three metabolites, although they also have reactions controlled by four metabolites. Table 1 summarizes the number of each type of interaction in each model. Because both biological models have significantly more metabolites and reactions than the synthetic models, the number of possible interactions that need to be considered is substantially greater.

### Autogenerated Training Data

Machine learning models must be trained using data that are broadly representative of the input data they are likely to encounter, which often entails using datasets that are as large as possible. In metabolism, there is a wide variety of metabolic reactions with disparate mechanisms and functional behaviors (e.g. bi-bi sequential reactions vs. ping-pong reactions) [25] or that are controlled by a different number of metabolites. However, appropriate training data for many of these possible situations are sometimes not available at all, let alone in sufficient quantity to enable machine learning model training. Accordingly, we chose to generate hundreds of artificial interactions to use as training data in an approach we refer to as “autogeneration”. While the practice of creating artificial training data has been used in other machine learning contexts before [26–29], to the best of our knowledge it has not been used in producing metabolite-dependent regulatory interaction data.

Damped sine waves with random parameters were generated to mimic common metabolite concentration profiles; these profiles were then used to calculate flux profiles using BST equations with random parameters [22]. Features for interactions labeled as true positives in the training data were calculated using these simulated concentration profiles and corresponding calculated flux profiles. The same process was used to calculate features for interactions labeled as true negatives, except that one or more of the input metabolite concentration profiles was replaced by a different, random metabolite concentration profile that was not used in the calculation of the target flux profile. This approach for autogenerating training data aims to circumvent the requirement for large dynamic metabolomics and fluxomics datasets to train the machine learning framework, which are currently not widely available on the scale that would be required. Because this autogeneration approach is independent of SCOUR, it can possibly be used for other computational methods that require an abundance of metabolic data. Further details on autogeneration of training data can be found in the Supplementary.

### Noisy Data

To generate noisy data that are more representative of what is expected to be acquired experimentally, we used two different sampling frequencies and two coefficients of variation (CoV) for randomly-added noise, for a total of four conditions. Sampling frequencies of 50 and 15 timepoints (nT) and CoVs of 0.05 and 0.15 were used, where a higher CoV represents more noise (experimental error). The smaller number of timepoints and the two levels of added noise are reasonable values for what one might expect from experimental mass spectrometry data for metabolomics or fluxomics. Starting with noiseless data, each metabolite and flux value in each time course was replaced with a random value drawn from *N_i,k_* ~ (*y_i_*(*t_k_*),*CoV*·*y_i_*(*t_k_*)), where *y_i_*(*t_k_*) is the value of species (metabolite or flux) *i* at time point *k*. For each timepoint, three noisy data values were generated to mimic triplicate samples, which is a common practice in metabolomics and fluxomics experiments.

### Data Pre-processing

For noisy data, we applied two different pre-processing steps to the data. For the one-controller metabolite interaction inference step of the framework, we used the median sample of the triplicate noisy data to calculate the features in the training and testing sets. For the two- and three-controller metabolite interaction inference steps, instead of using the medians, a moving Gaussian filter was applied to smooth the triplicate noisy data before calculating their features. The window size of the filter was chosen to be ¼ of the total simulation time, which was found to smooth the data without overfitting to the noise itself. While smoothing the noisy data for two- and three-controller metabolite interactions led to an increase in SCOUR’s performance, it was detrimental for one-controller metabolite interactions. We found that a few of the one-controller metabolite interaction features were more sensitive to the smoothed data than the noisy median data and would cause greater variability in SCOUR’s performance across repetitions.

### Features

Each step of the framework contains different “features” used to predict whether a particular interaction is likely to be correct. These “features” are scalar-valued outputs of functions that quantify characteristics of concentration and flux profiles and the relationships between different profiles, which may thus indicate whether a given metabolite or set of metabolites regulates a given flux. Different features were used for the prediction of interactions controlled by different numbers of metabolites (i.e., one-controller vs. two-controller vs. three-controller metabolite interactions). This allows the features to be customized to specific interaction types (and avoids the requirement that they must be valid or useful for all interaction types), which is expected to increase SCOUR’s overall accuracy compared to using the same features for all steps. Features were designed using biochemical insight into how metabolites are known to interact with enzymes and how these interactions would manifest in concentration and flux profile data. For example, for the one-controller metabolite interaction step, the Spearman correlation between fluxes and metabolites was used as a feature, as reaction fluxes are expected to be highly (though not necessarily linearly) correlated with the metabolites that control them. Additionally, features were created based on the expectation that for every set of metabolites that completely defines an output flux, each possible set of metabolite input concentrations can only yield one single output of flux reaction rate. A list and description of all features used in SCOUR can be found in Table S1.

### Machine Learning Algorithms and Stacking

Stacking is a technique used in machine learning that consolidates predictions from multiple machine learning algorithms [30]. Due to their different underlying assumptions and approaches, some machine learning approaches may identify regulatory interactions that other approaches miss. Combining predictions from different algorithms has been shown to boost overall accuracy. In SCOUR, we used four machine learning algorithms: random forest [31], k-nearest neighbors (kNN) [32, 33], shallow neural networks [34], and discriminant analysis [35]. Each of these algorithms are some of the most robust and commonly used machine learning approaches, but they are all fundamentally very different from one another. In SCOUR, we use kNN and discriminant analysis as binary classifiers: algorithms that can predict only two discrete labels. Random forest and neural networks are used as regression algorithms, where both predict continuous values. While most of these machine learning methods can be used as either discrete classifiers or regression models, we decided to use a mixture of these two types of algorithms because we believe that using a more diverse class of algorithms will minimize potential bias toward algorithms that are very similar to one another. In the stacking process, the predictions from the four models were used as the input of a secondary metamodel (another discriminant analysis classifier) to give a final classification output for each regulatory interaction that is tested. More details on the stacking method can be found in the Supplementary.

### Stepwise Approach

SCOUR uses a stepwise approach to identify different types of regulatory interactions at each step, beginning with the identification of one-controller metabolite interactions. First, two training datasets that consist of true positive (controlled by a single metabolite) and true negative (controlled by multiple metabolites) interactions are autogenerated for the two levels of the stacking model. The features described in Table S1 are calculated for each interaction in the first autogenerated dataset, which are used to train the random forest, kNN, shallow neural network, and discriminant analysis models in the first level of the stacking model. Next, the discriminant analysis model in the second level of the stacking model is trained using the feature matrix calculated from the second autogenerated dataset. Finally, the completely trained stacking model predicts whether each interaction in the testing dataset (comprising all possible one-controller metabolite interactions in the system of interest) is controlled by a single metabolite. This process is repeated for predicting two- and three-controller metabolite interactions.

This stepwise approach has two key advantages. First, it allows for completely independent classification models and features that can be crafted for specifically identifying reactions that are controlled by one, two, or three metabolites. We found (data not shown) that developing a one-step platform for simultaneous multiclass prediction (i.e. reactions controlled by different numbers of metabolites) led to worse performance even when using machine learning algorithms such as random forest and neural networks that can be tailored toward multiclass classification. Because SCOUR is trying to predict not just the number of metabolites controlling a reaction but also the exact controller metabolites, this additional layer of complexity is easier to address with multiple binary classification models.

The second advantage of using a stepwise approach is that after each step, fluxes whose regulatory status has already been identified are removed from consideration in the next step so that there are fewer interactions to be tested by the machine learning algorithms. This reduces the computation time of the entire stepwise process, reduces the chances of false positives, and allows subsequent steps and features to be more simply designed under the assumption that lower-order regulatory interactions will not be present in later steps. However, these advantages are at the risk of removing fluxes at a step earlier than when their true regulatory status could be identified. A comparison of the number of interactions that need to be tested whether or not the stepwise framework is used for each of the evaluated models can be found in Table 1. A schematic of SCOUR’s workflow is shown in Fig. 2.

**Fig. 2.**
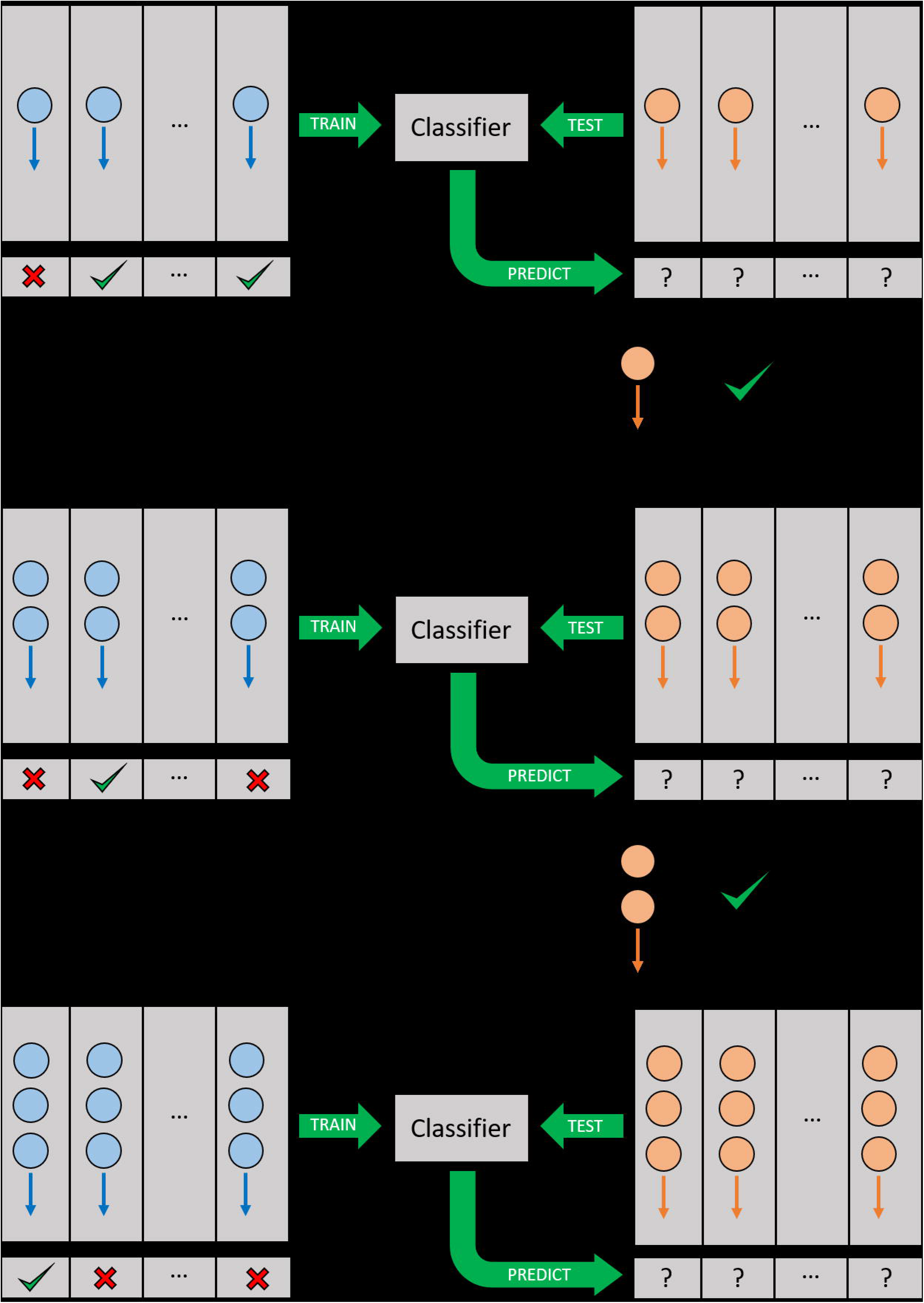
Workflow of stepwise machine learning framework for identifying one-, two-, and three-controller metabolite interactions. Blue circles and arrows represent metabolites and the fluxes they might interact with, respectively, in the training set. Orange circles and arrows represent metabolites and the fluxes they might interact with, respectively, in the testing set. In each step, the training set is used to train the machine learning classifier for fluxes with a specific number of metabolite controllers, which is then applied to the testing set to predict which fluxes are in that category. Between each step of the workflow, fluxes that have been positively identified are removed from further consideration in the test set.

### Framework Performance Metrics

To assess the performance of our framework in identifying different types of regulatory interactions, we evaluated four different metrics: accuracy, sensitivity, specificity, and positive predictive value (PPV). Accuracy is the percent of candidate regulatory interactions that are identified correctly as existing or not existing in a model. While accuracy can be a good metric if the classes (i.e. candidate interactions that are truly in the model vs. candidate interactions that are not in the model) are well-balanced, this is not the case for combinatorial consideration of potential metabolic regulatory interactions: there are many more candidate metabolite and reaction flux combinations than there are actual regulatory interactions in a given biological system. Sensitivity and specificity separate accuracy into two metrics that measure, respectively, the percent of positives (i.e. true regulatory interactions) that are identified correctly and percent of negatives (i.e. candidate regulatory interactions that are not actually in the model) that are identified correctly. PPV is the percentage of interactions predicted by the model that are true positives, an important metric to consider when one plans on experimental validation of predictions because it indicates how much effort is typically required for the validation of every newly discovered interaction. Exceedingly low PPVs are undesirable for predictions that are difficult to experimentally test, including metabolite-dependent regulation of reaction rates, because they signify that a large number of predicted interactions must be tested with these difficult experimental methods in order to find any true, validated interactions.

## Results

### Performance on Noiseless Data

When evaluating SCOUR on noiseless data, we found good overall predictive accuracy for both synthetic models (Fig. 3A-3B). We trained SCOUR on 30 independent sets of noiseless autogenerated data to assess the sensitivity of the framework to different sets of autogenerated training data. The average sensitivities and specificities for all steps in SCOUR were above 88% for both models. PPVs were above 77% for predicted one- and two-controller metabolite interactions, and above 58% for predicted three-controller metabolite interactions.

**Fig. 3.**
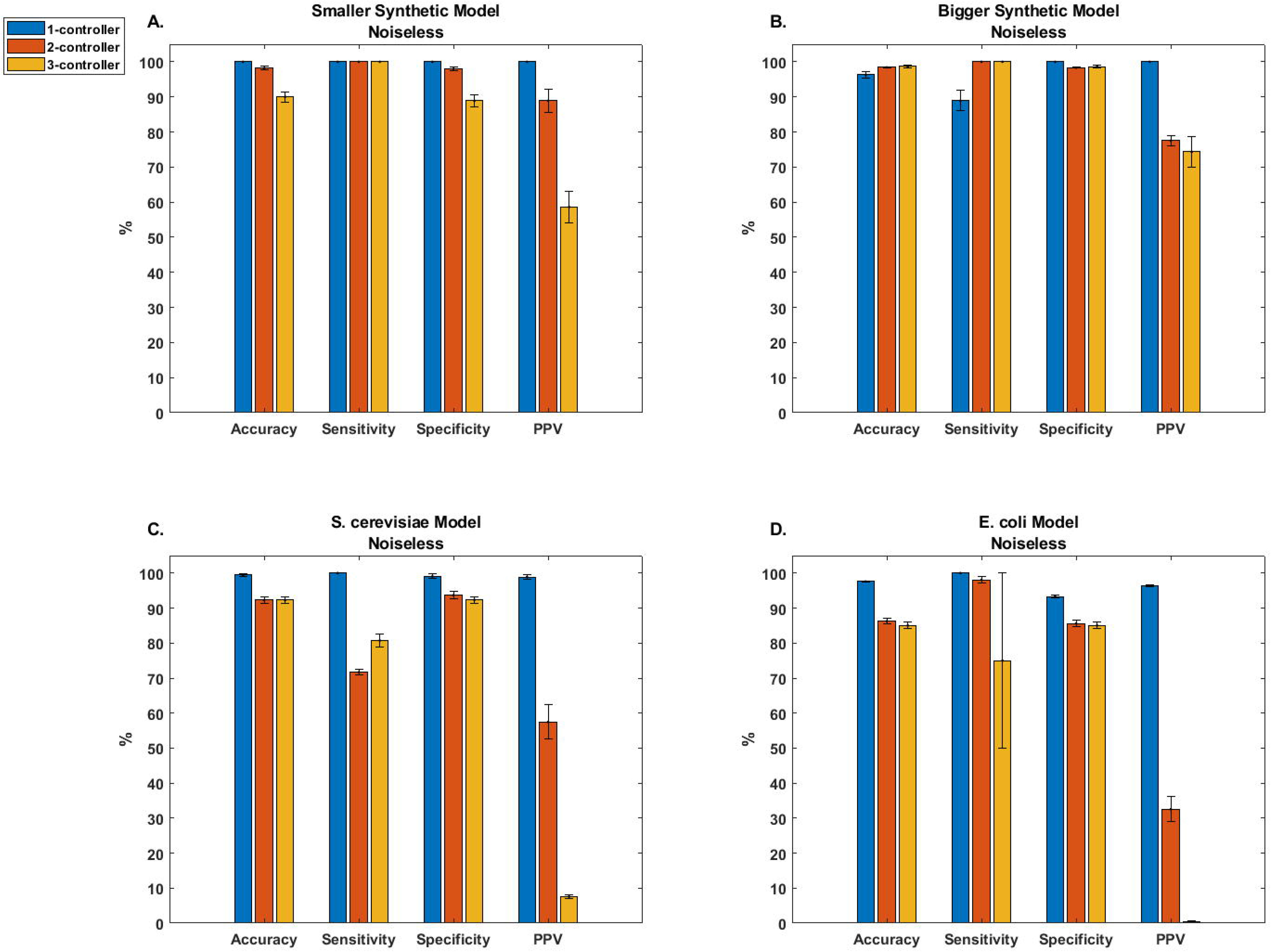
SCOUR performance on synthetic and biological models using noiseless training and test data. Bar graphs for accuracy, sensitivity, specificity, and PPV for each step of SCOUR in each tested model. Error bars represent the standard error of the mean (n = 30 from independent autogenerated training replicates).

We found similar results when testing SCOUR on noiseless data simulated from the *E. coli* and *S. cerevisiae* systems (Figure 3C-3D). As in the synthetic models, the PPV for both of these biological models decreased as the number of controller metabolites increased, though in a steeper fashion likely due to the increased complexity of these systems. The accuracy, sensitivity, and specificity in both biological models were still above 71% for all steps, and the PPVs for one- and two-controller metabolite interactions were all above 32%, despite the increase in model complexity. The low PPV for identification of three-controller metabolite interactions for both biological models (< 8%) despite high specificity (> 85%) was attributable to the highly imbalanced nature of the testing data. Out of the large number of candidate three-controller metabolite regulatory interactions that must be classified (Table 1), only a few are true positives and consequently there is an increased likelihood for false positive predictions. The large standard error of the mean for sensitivity in the *E. coli* model when identifying three-controller metabolite interactions is due to SCOUR removing the fluxes of the two true positive interactions in a previous step, which leads to the sensitivity not being calculated in several of the repetitions (due to the absence of any true positives or false negatives).

### Performance on Noisy Data

While using noiseless data gives a sense for the framework’s performance under ideal conditions, the realities of metabolomic and fluxomic experimental limitations can lead to significant deviation from these idealized assumptions. To assess SCOUR’s performance under more biologically relevant conditions, we examined two factors that need to be considered when using real metabolomics and fluxomics data: decreased experimental sampling frequency (and thus less information content to enable identification of true regulatory interactions) and increased experimental measurement noise. To give SCOUR a baseline performance level to compare to, we also created a classifier that randomly predicted whether a metabolite-flux interaction was a true positive or true negative interaction and used this to calculate a PPV at each step. Each interaction had a 50% chance of being classified as either a true positive or true negative in each step of the framework. For this random predictor, we assumed that the correct reaction fluxes were removed at each step, giving this classifier an advantage over our framework by greatly reducing the number of possible false positive interactions.

Assessment of SCOUR’s performance on noisy data from the synthetic models (Fig. 4A-4F) yielded similar trends to the results from noiseless data (Fig. 3A-3B). For both decreased sampling frequency and increased experimental noise (Fig. 4A-4F), SCOUR’s overall accuracy unsurprisingly decreased, but still allowed for effective identification of many regulatory interactions in each model. In both synthetic models, there was an expected decrease in sensitivity and PPV with decreasing sampling frequency or increasing noise. As in the noiseless case, the PPV decreased for fluxes with more controller metabolites due to the increase in candidate regulatory interactions (and thus, an increase in possible false positive predictions) tested at each stage. In the most experimentally realistic scenario (nT = 15, CoV = 0.15), SCOUR still yielded PPVs that were acceptable for lab validation when classifying one- and two-controller metabolite interactions (> 59% and > 18%, respectively). The mean PPVs for one- and two-controller metabolite interactions were also better than the random predictor in both synthetic models across all conditions, and SCOUR outperformed the random predictor in most cases when classifying three-controller metabolite interactions.

**Fig. 4.**
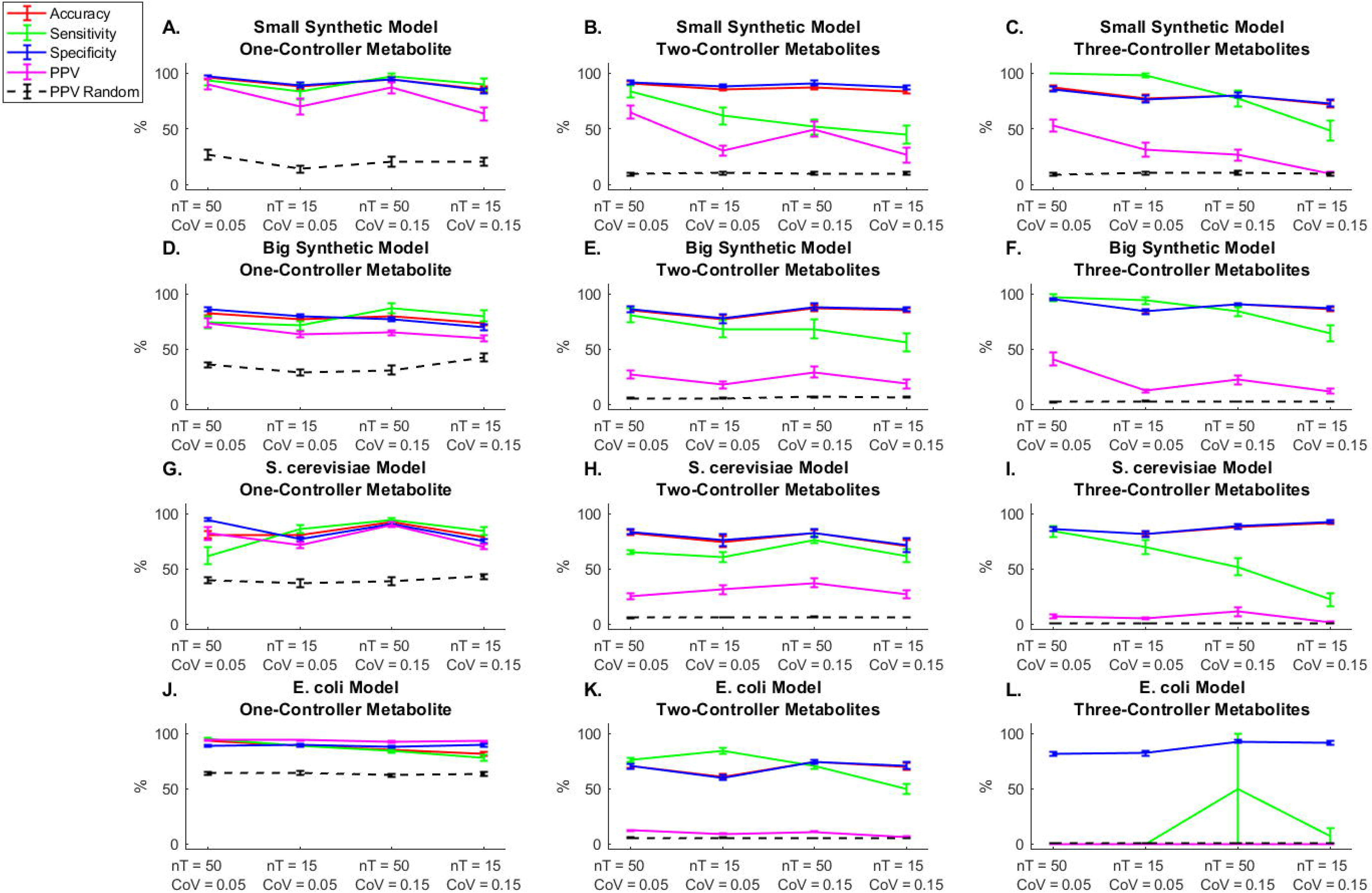
SCOUR performance on synthetic and biological models using noisy and low sampling frequency training and test data. Solid lines represent accuracy, sensitivity, specificty, and PPV performance of SCOUR on each model for each step of the framework. Dashed lines represent the PPV if interactions were randomly classified. Error bars represent the standard error of the mean. (n = 30 from independent autogenerated training replicates).

The results from testing on biological models with noisy data (Fig. 4G-4L) were similar to those from the synthetic models (Fig. 4A-4F). For both the *S. cerevisiae* and *E. coli* models, the PPV was fairly consistent (with slight decreases) for any given interaction type across the increasingly challenging noisy conditions, while accuracy, sensitivity, and specificity sometimes exhibited slightly more variability across those conditions. For the *S. cerevisiae* model, the PPV remained high (> 69% on average) in all conditions for identification of one-controller metabolite interactions and was above 25% for identification of two-controller metabolite interactions. These PPVs for one- and two-controller metabolite interactions are sufficiently high to allow for feasible experimental validation of these predictions to identify previously unknown interactions. For three-controller metabolite interactions, the PPV was below 12% for all conditions, which is not ideal from the standpoint of experimental practicality. In the *E. coli* model, the accuracy, sensitivity, specificity, and PPV were high for one-controller metabolite interactions in all conditions (> 92%), but the PPV dropped to less than 13% for two-controller metabolite interactions and was essentially 0% for three-controller metabolites (and a large standard error of the mean for sensitivity was observed, as in the noiseless condition in Fig. 3D). This would make it challenging to experimentally validate the *E. coli* predictions for two- and three-controller metabolite interactions without a guided high-throughput approach. Nevertheless, the PPVs for both biological systems when using SCOUR were on average better than the PPVs of the random predictor for one- and two-controller metabolite interactions for all conditions.

## Discussion

Our results indicate that SCOUR is a promising route for *in silico* prediction of metabolite-dependent regulation of metabolic fluxes using metabolomics and fluxomics data. On noiseless data, SCOUR predicts one- and two-controller regulatory interactions with high PPV in both synthetic and biological models, with three-controller interactions also predicted extremely well in some systems. While the use of noisy data leads to an expected drop in performance, SCOUR still provides extremely high PPV for one-controller interactions in all systems and high (experimentally useful) PPV for the synthetic models and the *S. cerevisiae* model when predicting two-controller metabolite interactions. SCOUR’s PPVs for these two steps greatly outperformed the PPVs of a random classifier in almost all cases. PPVs for three-controller metabolite interactions remained useful for the synthetic models but pushed the bounds of practical utility in the *S. cerevisiae* and *E. coli* models, likely attributable in large part to the combinatorial growth of the number of candidate interactions that must be tested and thus the concomitant growth in the number of false positives. Regardless of whether the three-controller interaction predictions are sufficient for experimental validation, the PPV in the *S. cerevisiae* model is significantly greater than the PPV for random classification for all noisy conditions except for the lowest sampling frequency and highest noise case (Fig. S2). This suggests that SCOUR would still be helpful for identifying these types of interactions compared to indiscriminate testing all combinations of interactions as higher-throughput guided experimental approaches are developed.

We believe the unusually sharp decrease in PPV for the *E. coli* model between the one- and two-controller interaction predictions is largely attributable to two reasons. First, the *E. coli* model contains three two-controller metabolite interactions where the two controller metabolites are highly correlated with each other (> 99% correlation; Fig. S3). This presents an identifiability problem, with it being extremely difficult to decouple the effects of the two metabolites once even a small amount of noise is added to their data. This in turn affected the utility of several features in our machine learning models, which led to these three interactions rarely being identified by SCOUR and thus also led to lower sensitivities and PPVs. Second, as previously discussed, the large size of the *E. coli* model necessitates testing many candidate regulatory interactions. Even with relatively high specificity, the resulting false positives from these tests can suppress the PPV. This is a common problem found in other efforts to determine regulatory activity (or any work with imbalanced datasets), where one class (e.g. true negative interactions) significantly outnumbers the other class (e.g. true positive interactions) [20].

Throughout the evaluation of SCOUR, we have relied on PPV as a performance metric because it is a valuable indicator of whether the predictions by SCOUR are worth experimentally validating. F1 score is another performance metric that is calculated from PPV and sensitivity and is often used for imbalanced datasets such as those found in this work. When using F1 scores, we found that SCOUR still outperformed random classification of one- and two-controller metabolite interactions in all models under all noisy conditions evaluated (Fig. S5). For three-controller metabolite interactions, the difference in performance between SCOUR and random classification is less clear and we would once again conclude that it would be difficult to recommend lab validation of the predictions for these types of interactions without high-throughput guided methods. While F1 score is an important evaluation metric and still verifies that SCOUR is a useful platform for identifying one- and two-controller metabolite interactions, we believe that PPV is a more important criterion because of its relevance to experimental validation: PPV indicates how many undiscovered interactions could be identified out of those predicted by SCOUR, regardless of how many true positive interactions exist.

Perhaps the most striking feature of SCOUR is its use of the autogeneration of synthetic interactions for training data. Because machine learning models generally require large amounts of data for training, and because this scale of data is typically not available for metabolomics and fluxomics data, we created a method to automatically generate training data that are in some way representative of a wide variety of real biological interactions. While these autogenerated “interactions” may not perfectly recapitulate the data that result from real reactions, SCOUR’s success shows that this autogeneration method can sufficiently train machine learning algorithms to identify regulatory interactions in many different systems. Because dynamic metabolomics and especially fluxomics data are so expensive and difficult to acquire with current analytical tools, this autogeneration method may prove useful for other tasks that require large amounts of these types of data.

Although this proof-of-principle framework has demonstrated significant potential for identification of many different regulatory interactions, there are several potential future avenues to improve overall performance. We note that both training and testing on autogenerated data produces higher PPVs (Fig. S6), which indicates that the autogenerated data do not perfectly capture biological interactions. Autogeneration of training data using Michaelis-Menten or other kinetics equations instead of BST equations could improve machine learning performance by generating training data that are more representative of the types of kinetics encountered in biological systems. While we chose to initially use BST equations for autogeneration based on their simplicity and utility in modeling many different systems [36], there are undoubtedly limitations to their generalization.

Second, as in all machine learning approaches, there is room to improve the features used to help predict true interactions. We designed knowledge-driven features based on how metabolites interact with the reaction fluxes they control. Data-driven features derived from raw metabolomics and fluxomics data could be beneficial if there are sufficient data to drive the derivation of these features, including the underlying “ground truth” about whether a given interaction truly exists in a system. Such features could include graph theoretical characterization of the network topology of how metabolites and fluxes are connected to each other, which has previously been used in metabolic contexts [37, 38]. However, an outstanding challenge will be how to include autogenerated data that are representative of these topological trends and capture the biological intricacies of metabolic systems, given that the autogenerated data are by definition synthetic and at least partly non-biological in nature. Additionally, machine learning algorithms for input to the stacking model beyond those tested here could also improve SCOUR’s performance.

Finally, the preprocessing of noisy experimental data undoubtedly can impact downstream analytical performance. While we settled on the median sample of the triplicate data when calculating features for one-controller metabolite interactions, and a Gaussian moving filter to smooth the data for two- and three-controller metabolite interactions, we also tried an average moving filter as well as an in-house smoothing approach [39]. Because the framework mostly produces extremely accurate results on noiseless data for all models tested, an improved data pre-processing approach (e.g., filtering, normalization, scaling, or other smoothing methods [40–42]) could significantly increase classification performance.

Notwithstanding these potential avenues for improvement, SCOUR is already a useful tool. At the very least, SCOUR can determine with high confidence reaction fluxes that are only controlled by a single metabolite, eliminating swaths of the metabolic network where metabolite-dependent regulation is unlikely to occur. However, SCOUR can also identify many more complex interactions, including possibly pointing towards reaction fluxes controlled by four or more metabolites (Fig. S4).

## Conclusions

SCOUR is a proof-of-principle for how metabolomics and fluxomics data can be leveraged with machine learning to find metabolite-dependent regulatory interactions; to our knowledge, this is the first reported example of such an approach. The identification of metabolite-dependent regulatory interactions has to date been critically hampered by experimental limitations in measuring and validating these interactions, making SCOUR’s predictions and triaging particularly valuable for such labor-intensive endeavors. Enabled by a method for autogenerating training data that reasonably mimic data from real biological systems, SCOUR circumvents the requirement for massive training sets that is typically associated with machine learning approaches. While metabolomics and fluxomics data are often collected at putative steady states, it is quite feasible to collect these data dynamically to leverage SCOUR’s potential for biological discovery. This means that as analytical methods for measuring metabolomics and fluxomics become cheaper and easier, and more data are available for analysis, SCOUR will be ready to take full advantage of these new datasets to discover biochemical regulatory interactions.

## Abbreviations

BST: Biochemical Systems Theory
CoV: Coefficient of Variance
nT: Number of timepoints
PPV: Positive Predictive Value
SCOUR: Stepwise Classification Of Unknown Regulation

## Declarations

### Ethics approval and consent to participate

Not applicable

### Consent for publication

Not applicable

### Availability of data and materials

The code created and used during the current study is available at http://github.com/gtStyLab/SCOUR

### Additional files

Additional file 1: Supplementary Information. Contains supplementary methods, tables, and figures. (PDF)

### Authors’ Contributions

JYL conceived of the study, participated in the design of the study, carried out computational experiments, analyzed experimental results, and helped to draft the manuscript. BN and CO carried out computational experiments and analyzed experimental results. MPS conceived of the study, participated in the design of the study, analyzed experimental results, and helped to draft the manuscript. All authors read and approved the final manuscript.

## Acknowledgements

Not applicable.

