## Supplementary figures and information for "SCOUR: A stepwise machine learning framework for predicting metabolite-dependent regulatory interactions"

### Supplementary Information

**Autogenerating Training Data**

Arguably the most important element of machine learning is the data used to train the machine learning algorithms. Because the training data need to capture a wide variety of possible input data that the classifier could be tested on, large training datasets are often required to achieve high predictive accuracy from machine learning methods. Due to current technical limitations in acquiring metabolomics and fluxomics data that limit the size of possible training datasets, we have created a novel approach for generating training data that emulates the interaction between controller metabolites and target fluxes in a process we refer to as "autogeneration".

For the training datasets in each step (meaning for the 1-controller, 2-controller, and 3-controller metabolite interactions), we created 300 autogenerated interactions, each with 15 different initial conditions. The 300 interactions included a mixture of samples that mimicked true positive and true negative interactions. To create the time course data for each controller metabolite, concentration profiles were created from damped sine wave functions with randomized parameters. $x_i$ is the concentration of controller metabolite $i$, $t$ is the simulation time, and $A$, $\lambda$, $\omega$, $\varphi$, are the amplitude, decay constant, angular frequency, and phase angle of a damped sine wave:

$$x_i = A_i e^{-\lambda_i t}(\cos(\omega_i t + \varphi_i) + \sin(\omega_i t + \varphi_i))$$

These concentration profiles were then used as input into Biochemical System Theory (BST) equations to calculate dynamic flux profiles. BST is an ordinary differential equation-based modeling framework for metabolic systems that uses power-law kinetics and is generalizable to many types of metabolic reactions [1]. Each BST equation also was assigned randomized parameters. $v$ represents the target reaction flux (rate), $x_i$ is the concentration of controller metabolite $i$, $n$ is the number of controller metabolites that regulate the target flux, and $\alpha$ and $\beta$ are the randomly assigned BST parameters.

$$v = \alpha \prod_{i=1}^{n} {x_i}^{\beta_i}$$

After generating both controller metabolite and target flux data, interactions were randomly assigned as either true positives or true negatives. If the interaction was labeled as a true positive, the correct sets of simulated controller metabolite profiles and corresponding calculated target flux profiles were used when calculating machine learning model features. However, if the interaction was labeled as a true negative, another set of metabolite concentration time course profiles would be generated from new damped sine wave functions and be used with the original target flux data to calculate features. Because these new pseudo-controller metabolites were not used in the calculation of the target flux data, there should be minimal relationship between the metabolites and the target flux, yielding a "true negative" data point.

To emulate the percentage of true positive interactions in the models tested in this work, of the 300 interactions in each step, 40%, 5%, and 5% were randomly assigned as existing interactions in the one-, two-, and three-controller interaction inference steps, respectively. Changes to these percentages are expected to shift the sensitivity and specificity of the framework, so it is important to base the training percentages on what is expected to be seen in the testing data based on existing biochemical knowledge.

**Scaling of feature matrices**

When assessing SCOUR's performance on noiseless data, we found that the feature matrices did not require any scaling. However, scaling the training and testing feature matrices significantly improved SCOUR's performance on noisy datasets. Each feature in the feature matrices for two- and three-controller metabolites were scaled between 0 and 1. When scaling data in the one-controller metabolite feature matrices between 0 and 1, we found poor performance due to a high sensitivity to outliers in a few of the one-controller metabolite features. To solve this problem, we scaled the feature matrices for one-controller metabolite interactions so that the $20^{th}$ and $80^{th}$ percentiles of the data were scaled between 0 and 1, which diminished the effect of outliers on the machine learning algorithms. This technique is called robust scaling [2].

**Features**

**Table S1 List of features for each step of the framework**

| **1-controller metabolite interaction features** | | |
|---|---|---|
| **Feature name** | **Description** | **Reasoning for feature** |
| Correlation | Spearman correlation between controller metabolite and target flux. | If a reaction flux is controlled by a single metabolite (most likely a mass action interaction), the Spearman correlation should be close to +1. |
| Curve fit | A second-degree polynomial is fit to the controller metabolite vs. target flux data and the adjusted $R^2$ value (adjusted for the number of coefficients in the polynomial model) is calculated. | A simple polynomial curve should fit the data reasonably well if a reaction is controlled by a single metabolite (e.g. if the data exhibits a Michaelis-Menten saturation curve). |
| Flux prediction | 2/3 of the available controller metabolite data and target flux data are randomly selected to train a kNN regression model, with the flux data acting as the dependent variable. The remaining 1/3 of controller metabolite data is used with the kNN model to predict the remaining flux values and the prediction error is calculated. This process is repeated 3 times and the mean of the prediction errors is taken. | It is likely easier to make predictions (i.e. lower prediction error) if the controller metabolite and target flux belong to an existing interaction in the system and no other metabolites truly regulate the target flux. |
| CoV of data | The average CoV of the target flux is calculated at 10 evenly-spaced individual concentrations within the range of the controller metabolite. Because flux data may not be sampled at these evenly-spaced concentrations, flux data are linearly | The CoV of the target flux should be low at each metabolite concentration for a one-controller metabolite interaction because there should be a single flux value for every concentration value. |

| | | |
|---|---|---|
| | interpolated at these concentrations using the closest higher and lower concentrations with sampled flux data. | |
| **2-controller metabolite interaction features** | | |
| **Feature name** | **Description** | **Reasoning for feature** |
| Functionality | Plot the two putative controller metabolites against each other for all 15 datasets produced from different initial conditions. Identify where the two controller metabolite concentrations are approximately equal to each other in two of the datasets by finding where the two datasets intersect with each other on the plot. This is accomplished by using the InterX function in MATLAB [3] that uses vectorization to determine intersection points between datasets. At these intersection points, linearly interpolate the flux data for each of the two datasets (using the scatteredInterpolant function in MATLAB for 3-D interpolation), calculate the difference of these two interpolated target flux values, and divide by the mean of the flux values to normalize. For all intersection points found, take the mean of all normalized differences between interpolated target fluxes. | For every input of controller metabolites, there should be a single output for the target flux (this is the definition of a mathematical function) if those metabolites are the only variables that interact with the reaction. |
| Surface fit | Fit a plane surface to the data of the two controller metabolite concentrations and the target flux and calculate the root mean square error of the fit against the data. | Because an existing interaction must maintain “functionality,” the controller metabolite and target flux data should form some sort of surface. There will likely be a better fit when fitting a plane to data from an existing interaction than data from a non-existing interaction. |
| Flux prediction | Same as flux prediction feature for 1-controller metabolite interactions, except two metabolites are used to train and test the kNN model. | Same as flux prediction feature for 1-controller metabolite interactions. |
| Correlation with one metabolite constant | Plot one of the putative controller metabolites (x-axis) against the target flux (y-axis) for each of the fifteen datasets. Next, plot ten vertical lines that are evenly-spaced within the range of the controller metabolite that represent ten constant concentrations. For one vertical line, identify if and where the line intersects with the fifteen data sets using the InterX function and linearly interpolate flux data at these intersection points using the closest higher and lower concentrations with sampled flux data. Calculate the Spearman correlation between the second controller metabolite and interpolated target flux at these intersection points where the first controller metabolite is constant. Repeat for each of the ten vertical lines and calculate the mean of all correlations. Switch which | The correlation between one controller metabolite and target flux should be consistently close to +1 (activation) or -1 (inhibition) for any constant concentration value for the second metabolite. This assumes no high concentration effects, such as substrate inhibition. The lesser of the two absolute mean correlations is taken as it is the worst performing. |

| | | |
|---|---|---|
| | metabolite is held constant and repeat the process. Take the lesser of the absolute values of the two mean correlations. | |
| Curve fit with one metabolite constant | Plot one of the putative controller metabolites (x-axis) against the target flux (y-axis) for each of the fifteen datasets. Next, plot ten vertical lines that are evenly-spaced within the range of the controller metabolite that represent ten constant concentrations. For one vertical line, identify if and where the line intersects with the fifteen data sets using the InterX function and linearly interpolate flux data at these intersection points using the closest higher and lower concentrations with sampled flux data. Fit a second-order polynomial to the second controller metabolite and target flux data at these intersection points and calculate the root mean square error between the fit and the data. Repeat for each of the ten vertical lines and calculate the mean of all errors. Switch which metabolite is held constant and repeat the process. Take the greater of the absolute values of the two mean errors. | If one controller metabolite is constant, a simple polynomial on the second controller metabolite and target flux should fit well, similar to the curve fit feature for 1-controller metabolite interactions. The greater of the two absolute mean errors is taken as it is the worst performing. |
| **3-controller metabolite interaction features** | | |
| **Feature name** | **Description** | **Reasoning for feature** |
| Hyperplane fit | Fit a hyperplane to the data of the three controller metabolite concentrations and the target flux and calculate the root mean square error of the fit against the data. | Same as surface fit feature for 2-controller metabolite interactions. |
| Functionality (percentage method) | For any two datasets generated from different initial conditions, find concentrations where the three putative controller metabolites are within 5% (noiseless) or 10% (noisy) across datasets. Calculate the CoV of the target flux for the two datasets at these points. | Same as functionality feature for 2-controller metabolite interactions. |
| Flux prediction | Same as flux prediction feature for 1-controller metabolite interactions, except three metabolites are used to train and test the kNN model. | Same as flux prediction feature for 1-controller metabolite interactions. |
| Functionality (rounding method) | For any two datasets generated from different initial conditions, find concentrations where the three putative controller metabolites are equal across datasets after rounding to the second decimal place. Calculate the percent of target flux values that are equal (within 0.002 error). The majority of metabolites in this work had mean concentrations on the order of $10^{-1}$ to $10^{1}$, and the majority of fluxes had mean rates on the order of $10^{-2}$ to $10^{2}$, making the chosen rounding precision | Same as functionality feature for 2-controller metabolite interactions. In this feature, the parameters used to determine equivalence (i.e. rounding precision and error threshold) are fixed and are not proportional to the concentration or flux data used, unlike in the percentage method. This difference allows the rounding method to be more sensitive when determining equivalence for concentration or flux data that have larger orders of magnitude (i.e. greater concentrations and fluxes will be considered equal in fewer |

| | | |
|---|---|---|
| | and error threshold reasonable for this feature. | cases than in the percentage method; if they are considered equal, it will be with higher confidence). The number of decimal places and error bounds can be adjusted depending on the user's preference for sensitivity. |
| Correlation with two metabolites constant | Same as correlation with one metabolite constant, except two metabolites are held constant and the Spearman correlation between the third metabolite and target flux is calculated. | Same as correlation with one metabolite constant. |

**Machine Learning Stacking**

Stacking is a technique used in machine learning to aggregate predictions made by multiple classification or regression algorithms [4]. The idea behind stacking is that some algorithms will be able to classify certain samples better than others, such that by combining information from multiple algorithms one can more accurately classify samples overall.

To train the four machine learning algorithms in the first layer of the stacking process, an initial set of autogenerated data with known training labels was used for each algorithm. To train the metamodel in the second layer of the stacking process, a second set of autogenerated data was passed through the previously trained first layer models and the prediction outputs from the four original machine learning algorithms were used as inputs to train the metamodel, along with the known training labels of the second set of autogenerated data. For this work, we chose to use a discriminant analysis classifier as the metamodel, as it was shown to perform well in consolidating information from the four algorithms in the first layer. A workflow of the stacking process is shown in Fig. S1. Training and testing of the stacking model was performed on a single CPU core in an Intel Xeon Gold 6226 processor with 2.70 GHz on the PACE clusters at the Georgia Institute of Technology using MATLAB R2019a [5]. Each repetition took ~2-6 CPU-hours (dependent on the number of timepoints in the testing data) .

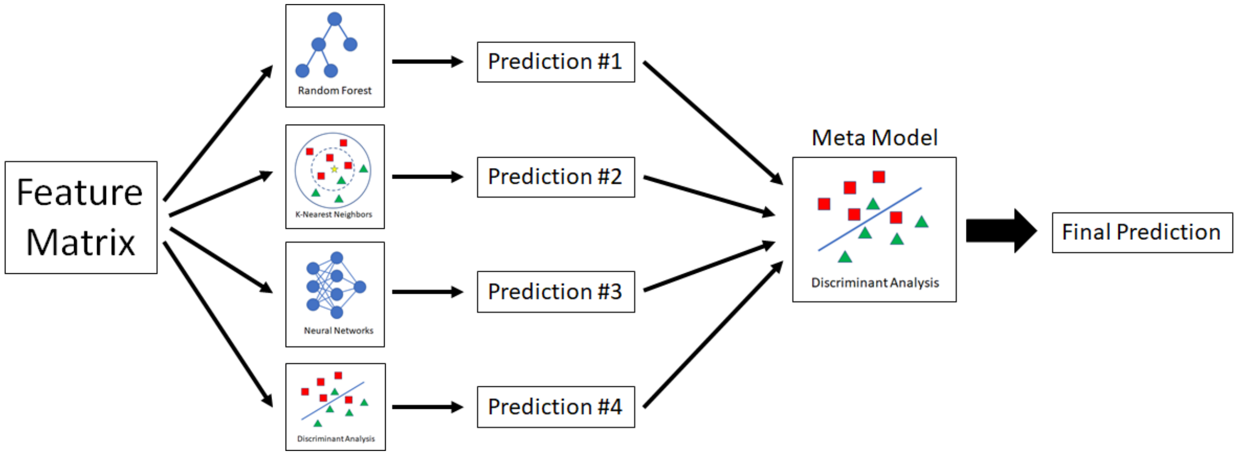


**Fig. S1. Workflow of stacking process.** After the feature matrix is calculated for all possible regulatory interactions, it is used on the first level of the stacking process as input for the four machine learning algorithms. The four resulting predictions are then used in the second level metamodel to produce a final prediction output for the framework.

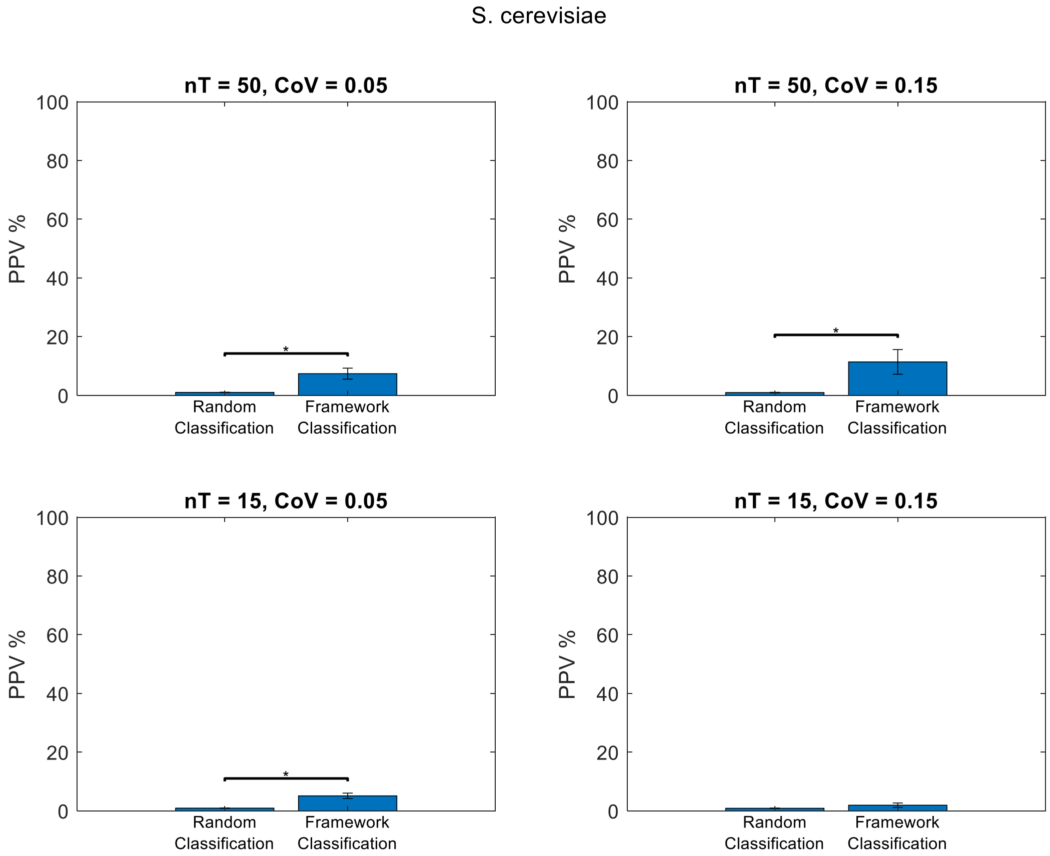


**Fig. S2. SCOUR's PPV for 3-controller metabolite interaction predictions is significantly greater than a random classifier.** While SCOUR yields low PPVs in the *S. cerevisiae* model for 3-controller metabolite interactions, these results are significantly better than random classification of 3-controller metabolite interactions for all conditions except for the case with the fewest time points and most noise (nT = 15, CoV = 0.15). We used a Wilcoxon rank-sum test (alpha = 0.05) to assess significance, as we found the distributions of the PPVs from SCOUR were not normal when using a Kolmogorov-Smirnov test (alpha = 0.05), except for the nT = 50, CoV = 0.05 condition. With appropriate guided high-throughput methods, SCOUR's predictions could still be useful for identifying these types of reactions.

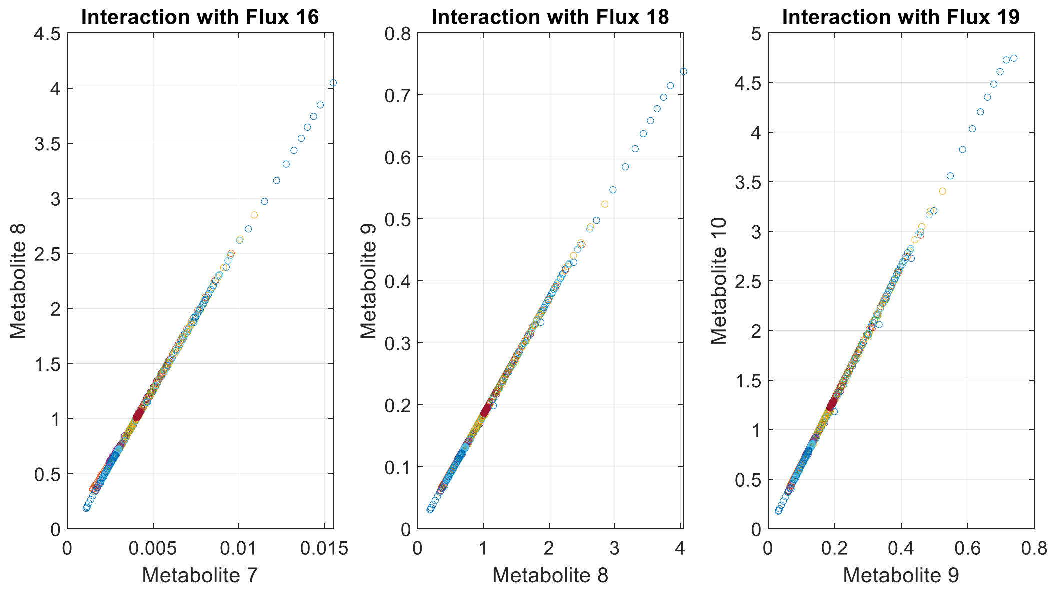


**Fig. S3. The most common false negative two-controller metabolite interactions in the *E. coli* model.** Three two-controller metabolite interactions were not correctly predicted across many conditions and replicates. The metabolites involved in these three interactions were highly correlated with each other, as indicated by these plots of noiseless concentration data for each of the pairs of metabolites. Each of the fifteen datasets with different initial conditions is represented by a different group of colored data. The scatter plots indicate extremely high correlation, which mitigated the utility of some of the features provided to the machine learning models (e.g. functionality and flux prediction). Inclusion of such interactions with highly correlated controller metabolites in the autogenerated data did not improve SCOUR's ability to identify these metabolites, especially once noise was introduced into the measurements.

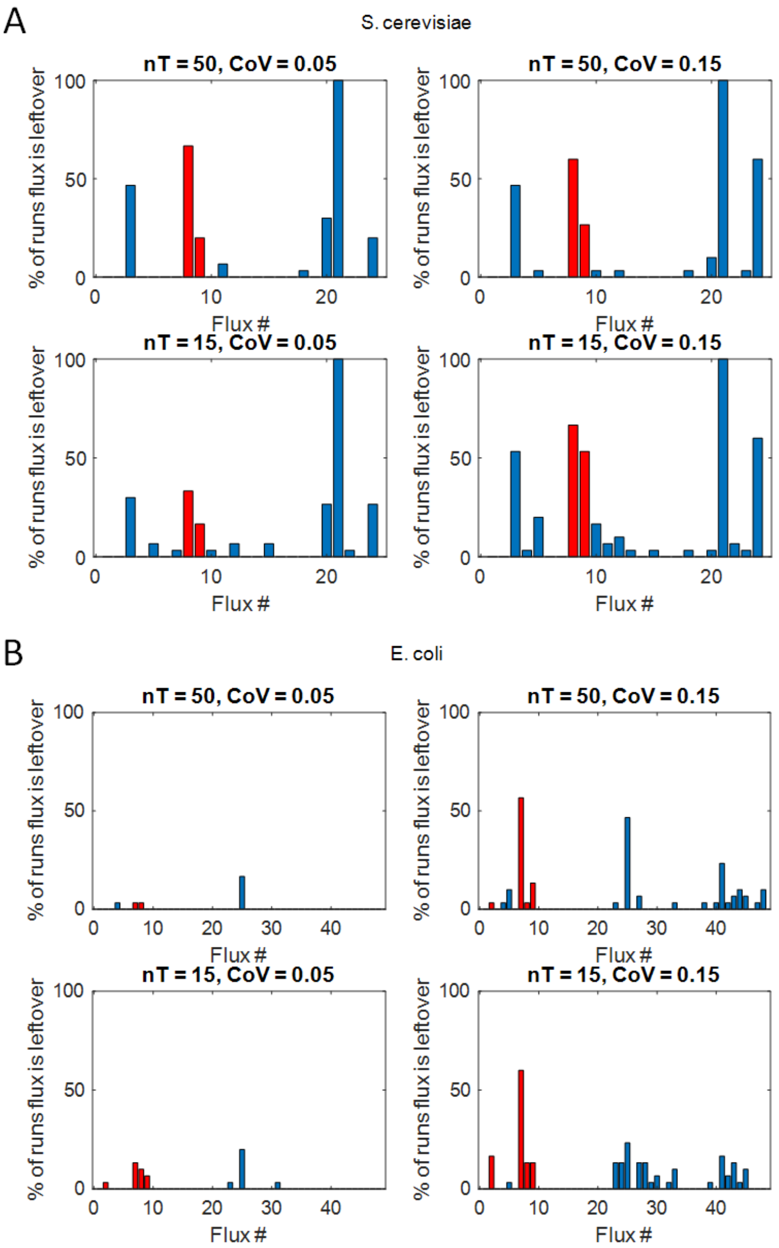


**Fig. S4. Four-controller and higher-order metabolite regulatory interactions.** While we only focus on classifying one-, two-, and three-controller metabolite interactions, the framework may still provide information about four-controller or higher-order metabolite interactions. After removing the fluxes with predicted regulatory relationships in the three described steps of the framework, we have found that there is some evidence supporting the identification of leftover fluxes in the A) *S. cerevisiae* and B) *E. coli* models as being controlled by four metabolites. The two models contain 2 and 4 four-controller metabolite interactions (red bars), respectively. While the framework is unable to identify the controller metabolites that interact with these fluxes, being able to predict which fluxes are controlled by more than three metabolites may be useful for understanding reaction mechanisms. Further improvements to the accuracy in each of the steps of the framework would be required to ensure that the majority of fluxes that remain are truly controlled by four or more metabolites.

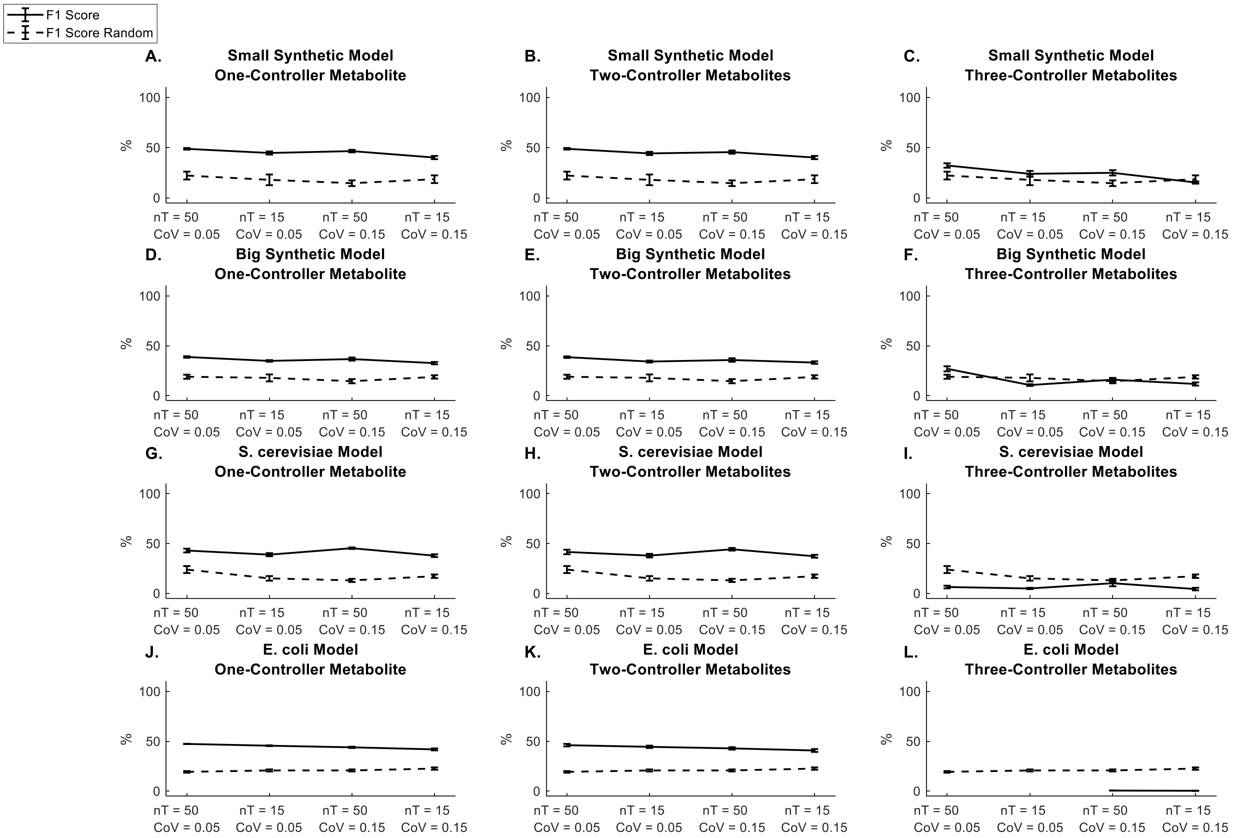


**Fig. S5. F1 scores for synthetic and biological models using noisy and low sampling frequency training and test data.** Bold lines represent the average F1 scores of SCOUR and dashed lines represent the average F1 scores when randomly classifying interactions (n = 30 from independent autogenerated training replicates). Note that the F1 score does not exist when identifying three-controller metabolites in the *E. coli* model for two conditions (nT = 50, CoV = 0.05 and nT = 15, CoV = 0.05) because SCOUR had removed both of the true positive three-controller metabolite interactions in a previous step of the framework for all repetitions, meaning sensitivity (and therefore F1 score) could not be calculated.

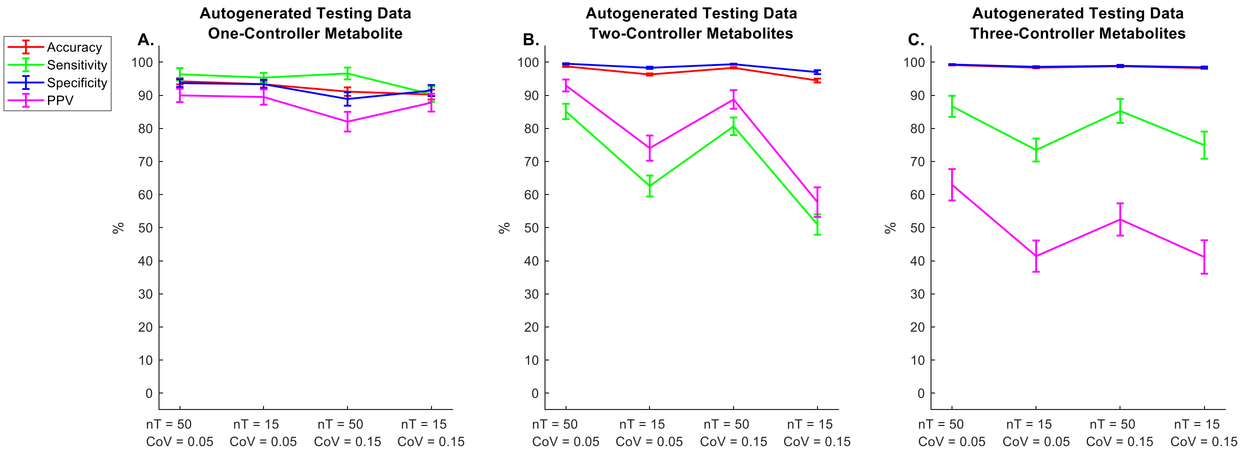


**Fig. S6. SCOUR performance when training on autogenerated data and testing on autogenerated testing data.** Testing data were autogenerated in a similar manner to the training data, but the percentage of true positive three-controller metabolite interactions in the testing set was set to 1% to more closely resemble the percentages found in the synthetic and biological models assessed (Note: training SCOUR with 1% instead of 5% true positives did not seem to improve performance). The results when testing on autogenerated data were better than when testing on the synthetic or biological models, which is likely due to the testing data more closely resembling the training data because both datasets were autogenerated. The fact that we do not see consistently high performance across all sets of the framework even for the training data, similar to when testing on the synthetic and biological systems, suggests that SCOUR is likely not overfitting.
